# XenoCP: Cloud-based BAM cleansing tool for RNA and DNA from Xenograft

**DOI:** 10.1101/843250

**Authors:** Michael Rusch, Liang Ding, Sasi Arunachalam, Andrew Thrasher, Hongjian Jin, Michael Macias, Lawryn Kasper, Andre Silveira, Michael A. Dyer, Suzanne J. Baker, Jinghui Zhang

## Abstract

**Summary:** Xenografts are important models for cancer research and the presence of mouse reads in xenograft next generation sequencing data can potentially confound interpretation of experimental results. We present an efficient, cloud-based BAM-to-BAM cleaning tool called XenoCP to remove mouse reads from xenograft BAM files. We show application of XenoCP in obtaining accurate gene expression quantification in RNA-seq and tumor heterogeneity in WGS of xenografts derived from brain and solid tumors.

**Availability and Implementation:** St. Jude Cloud (https://pecan.stjude.cloud/permalink/xenocp) and St. Jude Github (https://github.com/stjude/XenoCP)

## 1 INTRODUCTION

The patient derived xenograft (PDX) model is a powerful tool for understanding cancer development and for exploring new therapeutic treatment as these models are expected to faithfully recapture tumor growth, metastasis and disease outcomes^1,2^. In PDX models the host mouse cells admix with the patient tumor cells upon engraftment, which is known as mouse contamination. The presence of mouse contamination poses a challenge for Next Generation Sequencing (NGS) analysis, as it can confound the genomics analysis and interpretation of the data.

Many software programs are available to separate or remove the mouse reads in NGS analysis. These programs utilize either a k-mer method or quality/mapping scores^3,4^. Some programs work only with specific aligners^5^, give multiple bam outputs for selection by end user^6^, or no implementation is readily available^7^.

We developed a cloud-based tool called XenoCP that identifies and removes contaminating host reads in PDX NGS data by comparing existing graft genome alignments to realignment to the host genome (Fig1A). To our knowledge this is the first cloud-based tool designed to address xenograft contamination. XenoCP can be easily incorporated into any workflow as it takes a BAM file as input yields a clean BAM output, preserving existing alignments for non-contaminant reads. It can also be run as a containerized application, greatly simplifying the installation.

**Figure 1.**
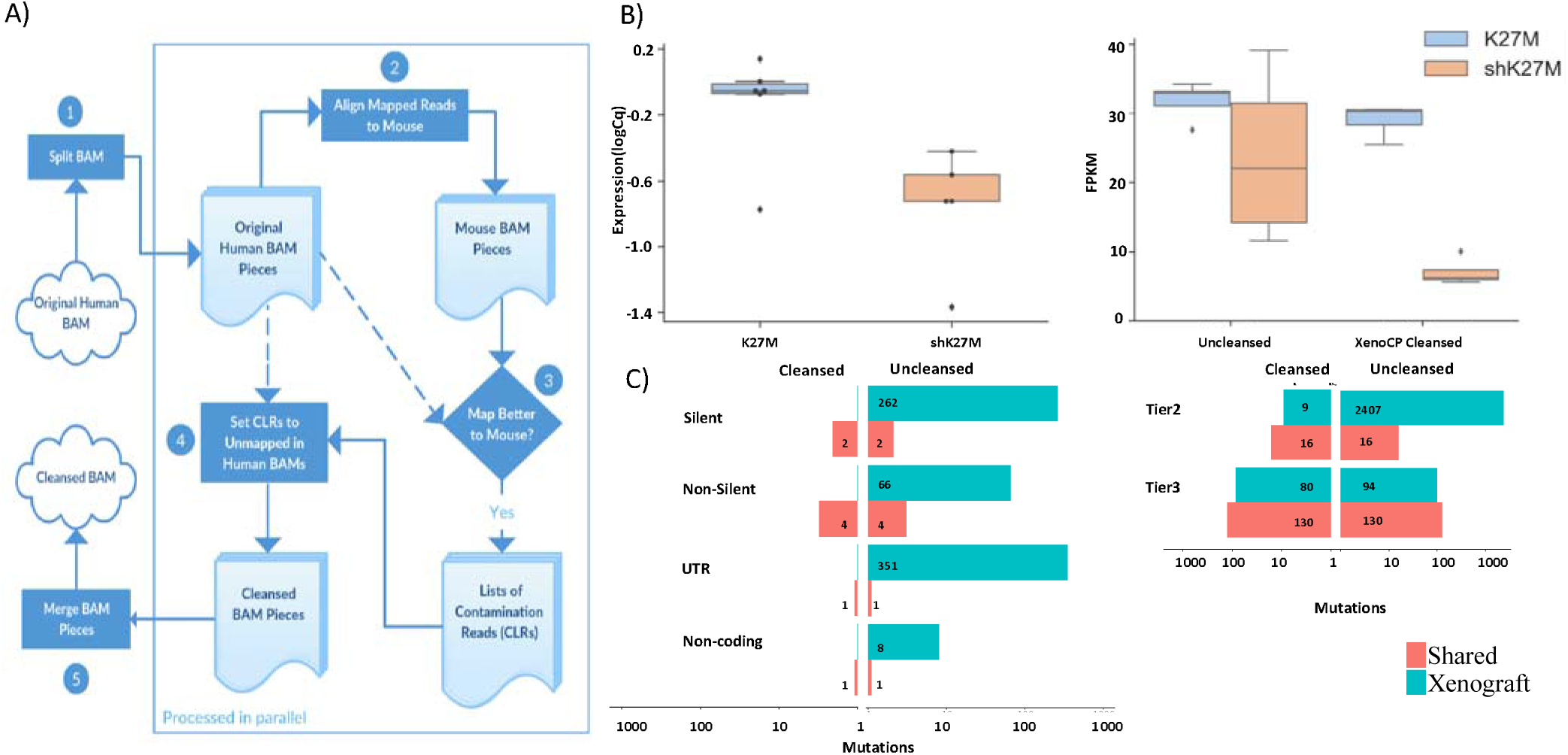
A) XenoCP algorithm. B) Gene quantification of xenograft samples. (Left) Potent knockdown of H3F3A (K27M) as measured by qRT-PCR. Values shown on plots are normalized by GAPDH. (Right) Comparisons of H3F3A FPKM values before and after XenoCP cleaning. C) Number of xenograft SNV calls shared with diagnosis or in xenograft only, before and after XenoCP cleaning. (Left) Tier 1 (silent, non-silent, UTR) and (Right) tier 2 (conserved) and tier 3 (non-repetitive) calls.

We tested XenoCP on simulated whole genome sequencing (WGS) data as well as RNA-Seq and WGS from xenograft samples. Removal of mouse reads facilitated accurate gene expression quantification in RNA-seq and removal of false positive somatic variants in WGS.

## 2 METHODS

XenoCP accepts an existing alignment to the graft (typically human) genome in BAM format. Aligned reads are re-aligned to the host (typically mouse) genome using the Burrows-Wheeler Aligner(BWA)^8^. Each mapping is scored using matching bases less gap count. If a read’s host mapping has a higher score than its graft mapping, then the read and its mate are both considered contamination. XenoCP outputs a copy of the original BAM with contamination reads marked as unmapped (Fig. 1A). Further details are described in Supplementary Methods.

## 3 RESULTS

First, we simulated a xenograft BAM file by mixing reads from human and mouse files (Supplementary Methods) and XenoCP achieved 86.40% sensitivity and 99.98% specificity on C56BL/6 (Supplementary Table 1).

Next, we used XenoCP to remove mouse reads from RNA-Seq. We knocked down the *H3F3A* (K27M) gene in a diffuse intrinsic pontine glioma (DIPG)^9^ xenograft sample (referred to as sh^K27M^) (Supplementary Methods). Quantitative real-time reverse transcriptase polymerase chain reaction (qRT-PCR) confirmed the robust knockdown of *H3F3A* relative to a control (K27M). Initial analysis of RNA-Seq generated from the samples was inconsistent with the qRT-PCR results. After cleaning with XenoCP, gene expression of *H3F3A* recapitulated the qRT-PCR result (Fig. 1B), demonstrating the necessity and effectiveness of XenoCP cleaning in the accurate gene quantification of xenograft samples.

Finally, we applied XenoCP to remove mouse reads from a PDX whole genome sequencing (WGS) sample. WGS reads for germline, primary diagnosis and xenograft from a retinoblastoma were aligned to GRCh37-lite using BWA. The xenograft alignment was cleansed using XenoCP. Somatic single nucleotide variants (SNVs) were called using Bambino^10^.

Somatic variants from diagnosis, uncleansed xenograft and cleansed xenograft were manually reviewed (Fig1C) and validated by targeted capture sequencing at an average of >350X coverage as described previously^11^. Using cleansed PDX data, we found that the majority of SNVs were shared between diagnosis and PDX. By contrast, using the uncleansed PDX data, the majority of variants were classified as Xenograft-specific (Fig1C). For example, tier1 mutations, defined as coding, UTR and non-coding RNA mutations, are identical in the patient sample and xenograft using the cleansed PDX data. By contrast, the uncleansed bam file produced an extra 687 tier1 mutations specific to the xenograft sample, showing that reads from the mouse genome in the uncleansed data could lead to misinterpretation regarding the genetic fidelity between patient tumor and xenograft mouse model (Fig. 1C). Results from the non-coding regions where tier2 (conserved regions) and tier3 (non-repetitive) mutations are located follow the same trend: the vast majority (2,391 out 2,481. 96%) of the PDX-specific mutations were caused by contaminating mouse reads which were removed after XenoCP cleansing.

In summary, we present XenoCP, a cloud-based BAM-to-BAM cleansing tool to remove mouse reads from DNA or RNA xenograft NGS data, increasing accuracy of gene expression and variant calling.

## ACKNOWLEDGMENTS

This project was funded in part by the American Lebanese Syrian Associated Charities (ALSAC). Research reported in this publication was supported in part by the Cancer Center Support grant P30 CA021765 from the National Cancer Institute of the National Institutes of Health and by Collaborating Mutations in Medulloblastoma grant P01 CA096832 from the National Cancer Institute of the National Institutes of Health.

## REFERENCES

1. Davies, A. H., Wang, Y. & Zoubeidi, A. Patient-derived xenografts: A platform for accelerating translational research in prostate cancer. Mol. Cell. Endocrinol. (2018). doi:10.1016/j.mce.2017.03.013

2. Yoko, S. D. et al. Tumor grafts derived from women with breast cancer authentically reflect tumor pathology, growth, metastasis and disease outcomes. Nat. Med. (2011). doi:10.1038/nm.2454

3. Conway, T. et al. Xenome-a tool for classifying reads from xenograft samples. Bioinformatics (2012). doi:10.1093/bioinformatics/bts236

4. Callari, M. et al. Computational approach to discriminate human and mouse sequences in patient-derived tumour xenografts. BMC Genomics (2018). doi:10.1186/s12864-017-4414-y

5. Ahdesmäki, M. J., Gray, S. R., Johnson, J. H. & Lai, Z. Disambiguate: An open-source application for disambiguating two species in next generation sequencing data from grafted samples. F1000Research (2017). doi:10.12688/f1000research.10082.2

6. Khandelwal, G. et al. Next-Generation Sequencing Analysis and Algorithms for PDX and CDX Models. Mol. Cancer Res. (2017). doi:10.1158/1541-7786.mcr-16-0431

7. Rossello, F. J. et al. Next-Generation Sequence Analysis of Cancer Xenograft Models. PLoS One (2013). doi:10.1371/journal.pone.0074432

8. Li, H. Aligning new-sequencing reads by BWA BWAL□: Burrows-Wheeler Aligner. Slides (2010).

9. Silveira, A. B. et al. H3.3 K27M depletion increases differentiation and extends latency of diffuse intrinsic pontine glioma growth in vivo. Acta Neuropathol. (2019). doi:10.1007/s00401-019-01975-4

10. Edmonson, M. N. et al. Bambino: A variant detector and alignment viewer for next-generation sequencing data in the SAM/BAM format. Bioinformatics (2011). doi:10.1093/bioinformatics/btr032

11. Zhang, J. et al. A novel retinoblastoma therapy from genomic and epigenetic analyses. Nature (2012). doi:10.1038/nature10733

