## supplementary for "XenoCP: Cloud-based BAM cleansing tool for RNA and DNA from Xenograft"

Supplementary Material for XenoCP

**Supplementary Methods**

**Detailed description of XenoCP**

The pipeline accepts an existing alignment to the graft (typically human) genome in BAM format. The input BAM is split into pieces, processed to mark contaminating reads as unread, and then reconstituted into a single purified BAM. For paired-end sequencing, the algorithm processes each mate pair as a single unit since mate reads are from the same physical insert. Therefore, the splitting step is designed to keep read pairs together, rather than being a naïve split by region or number of reads.

Mapped reads in each BAM piece are then analyzed to find contamination. Mapped reads are extracted from the BAM in FASTQ format and are then mapped to the host (typically mouse) reference genome using the Burrows-Wheeler Aligner (BWA). Each read’s original mapping is compared with the host mappings to call the read as from graft or host, as follows. Reads not mapping to any host assembly are called graft. Remaining reads’ original and host mappings are assigned a score which is the number of base matches less the number of gaps. If a read has a host mapping with a higher score than the original mapping, then both the read and its mate (if paired) are called as host contamination. Finally, reads called as host contamination are set to unmapped, and the updated BAM pieces are merged into the final cleansed BAM. Contamination read names and reads with identical original/host scores are preserved for further analysis.

**In silico validation by simulation**

We simulated murine contamination in a human xenograft sample by generating WGS BAM files that each consisted of 9% mouse reads (C57BL/6 or FVB) and 91% human reads. The human reads were subsampled from human retinoblastoma primary tumor WGS reads. Mouse reads were subsampled from either C56BL/6 (EBI acc. ERP000041) or FVB (EBI acc. ERP0000687). The pipeline was run using both MGSCv37 and CELERA assemblies of the mouse genome.

After running XenoCP, we counted the number of reads from the human sample an reads from the mouse sample that were either removed or retained by XenoCP and then calculated sensitivity (removed mouse reads divided by total mouse reads) and specificity (retained human reads divided by total human reads). We noted that especially in FVB many of the retained mouse reads in FVB contained the telomeric repeat sequence, so these numbers are indicated as well in Supplementary Table 1.

***H3F3A* Knockdown**

We knocked down the *H3F3A* (K27M) gene using short hairpin RNA (shRNA) in three diffuse intrinsic pontine glioma (DIPG)^9^ xenograft models (referred to as sh^K27M^). A non-silencing shRNA with no target in the human genome (referred to as K27M) was used as a control. Quantitative real-time reverse transcriptase polymerase chain reaction (qRT-PCR) confirmed the robust knockdown of *H3F3A.* However, the analysis of RNA-Seq generated from the paired xenograft samples for one of the three models showed inconsistency with the qRT-PCR results using the uncleansed bam file, and this sample was used for the presented analysis.

**Supplementary Data**

The supplementary data file contains SNV calls with read counts from primary sequencing and capture validation. Note that some variants had not been discovered as part of the original study and were therefore not included in capture validation. These have N/A in the validation read counts.

**Supplementary Table 1**

| Metric | C57BL6 | FVB |
| --- | --- | --- |
| Mapped Reads | 1,058,313,934 | 1,058,389,524 |
| Human | 1,058,084,926 | 1,058,084,926 |
| Removed | 223,340 | 223,368 |
| Retained | 1,057,861,586 | 1,057,861,558 |
| Mouse | 229,008 | 304,598 |
| Removed | 197,852 | 214,524 |
| Retained | 31,156 | 90,074 |
| Non-Telomeric | 22,729 | 24,246 |
| Sensitivity | 86.40% | 70.43% |
| Specificity | 99.98% | 99.98% |
